# Tunable human myocardium derived decellularized extracellular matrix for 3D bioprinting and cardiac tissue engineering

**DOI:** 10.1101/2021.03.30.437600

**Authors:** Gozde Basara, S. Gulberk Ozcebe, Bradley W. Ellis, Pinar Zorlutuna

**Affiliations:** Aerospace and Mechanical Engineering Department, University of Notre Dame, Notre Dame, Indiana 46556, USA; Bioengineering Graduate Program, University of Notre Dame, Notre Dame, 46556, IN, USA

**Keywords:** 3D bioprinting, decellularized extracellular matrix, cardiac tissue engineering, gelatin methacryolyl, methacrylated hyaluronic acid, human induced pluripotent stem cell-derived cardiomyocyte, human cardiac fibroblasts, dual crosslinking

## Abstract

The generation of 3D tissue constructs with multiple cell types and matching mechanical properties remains a challenge in cardiac tissue engineering. Recently, 3D bioprinting has become a powerful tool to achieve these goals. Decellularized extracellular matrix (dECM) is a common scaffold material due to providing a native biochemical environment. Unfortunately, dECM’s low mechanical stability prevents usage for bioprinting applications alone. In this study, we developed bioinks composed of decellularized human heart ECM (dhECM) with either gelatin methacryloyl (GelMA) or GelMA- methacrylated hyaluronic acid (MeHA) hydrogels dual crosslinked with UV light and microbial Transglutaminase (mTGase). We characterized the bioinks’ mechanical, rheological, swelling, printability and biocompatibility properties. Composite GelMA-MeHA-dhECM (GME) hydrogels demonstrated improved mechanical properties by an order of magnitude, compared to GelMA-dhECM (GE) hydrogels. All hydrogels were extrudable and compatible with human induced pluripotent stem cells derived cardiomyocytes (iCMs) and human cardiac fibroblasts (hCFs). Tissue-like beating of the printed constructs with striated sarcomeric alpha-actinin and Connexin 43 expression was observed. The order of magnitude difference between the elastic modulus of these hydrogel composites offers applications in *in vitro* modelling of the myocardial infarct boundary. Here, as a proof of concept, we created an infarct boundary region with control over mechanical properties along with cellular and macromolecular content through printing iCMs with GE bioink and hCFs with GME bioink.

## 1. Introduction

Myocardial infarction (MI) is one of the most common cardiovascular diseases and has remained the main cause of death worldwide for decades [1,2]. Although many studies are dedicated to understanding the MI mechanism and potential therapeutic options on animals, research concerning human myocardium remains very limited [3]. For this reason, it is crucial to fabricate human benchtop models recapitulating the mechanical properties of both healthy cardiac tissue and fibrotic scar tissue, which is formed as a result of the collagen deposition and fibroblast activation triggered in response to the MI and has an increased stiffness [4].

3D bioprinting is one of the most powerful tools used to fabricate mimetic cardiac tissues, capturing the complexity of the native cellular composition and matrix structure [5–8]. These printed constructs have been used in different applications, including but not limited to drug screening and regenerative medicine [5,7,8]. Extrusion based bioprinting is one of the most commonly used 3D bioprinting methods [9–12], in which cell-laden bioinks are extruded on a platform following a pre-designed shape. For this process, the viscoelastic properties of the bioinks are explicitly important for printability while ensuring cell viability [4]. Natural hydrogels like collagen [13,14], gelatin [15,12,16], alginate [17,18], and hyaluronic acid (HA) [19,20] have been widely used as bioinks for extrusion based bioprinting. Yet they fall short on fully capturing the biochemical cues of the native tissue microenvironment.

To obtain naturally derived biomaterials recapitulating the biochemical cues in the native tissue environment, decellularization techniques have been developed and decellularized ECM (dECM) has become a preferred biomaterial for cardiac tissue engineering applications such as cardiac repair and regeneration [21–25]. Although dECM preserves the biochemical composition of the native tissue, it loses the mechanical and structural stability of tissue during the decellularization and digestion processes. Therefore, for printing purposes, dECM is usually crosslinked via mixing it with B2 vitamin [24], adjusting the pH [22], or combined with some other natural or synthetic biomaterials [4,13,21,25–30]. These biomaterials provide additional control over scaffold stiffness which can be used to mimic the parameters of cardiac tissues of varying ages or pathologies.

Gelatin methacryolyl (GelMA) is a very commonly used natural hydrogel for both cardiac tissue engineering and 3D printing applications [12,31,32]. Methacrylated HA (MeHA) has been commonly used in bone tissue engineering applications due to its increased rigidity and resistance to degradation [33]. Previous studies have shown that GelMA-MeHA hybrid hydrogels display highly tunable physical properties and can be used for different applications [19,34]. Unfortunately, the literature lacks a complete mechanical characterization, compatibility, and printability investigation of GelMA and GelMA-MeHA hydrogels mixed with dECM for cardiac tissue engineering applications.

In our study, we blended GelMA and GelMA-MeHA hydrogels with decellularized human cardiac ECM (dhECM) to create cardiac tissue-like constructs. We measured the elastic modulus of GelMA (G), GelMA-dhECM (GE), GelMA-MeHA (GM) and GelMA-MeHA-dhECM (GME) hydrogels which were dual crosslinked using UV and microbial Transglutaminase (mTGase) and observed an order of magnitude difference between GE and GME hydrogels. The fabricated gels were then characterized in terms of rheology, swelling, printability, and compatibility of human induced pluripotent stem cell-derived cardiomyocytes (iCMs) and human cardiac fibroblasts (hCFs). We then characterized iCMs printed with G and GE in terms of beating properties and protein expression. Lastly, knowing that there is an approximately ten-fold difference between the stiffness of healthy cardiac tissue (8-12 kPa) and scar tissue (>150 kPa) formed after myocardial infarction (MI), an infarct region model was printed using a dual printhead by mixing iCMs with GE to model the healthy tissue and hCFs with GME to represent the scar tissue. To our knowledge, this is the first study to characterize GE, GME composite hydrogels and demonstrate their potential through the printing of an infarct boundary zone using iCMs and hCFs to create a human myocardial infarct boundary model.

## 2. Results and Discussion

### 2.1. Decellularization and Solubilization of Human Cardiac Tissue

Hearts that were deemed unsuitable for transplant were removed by the acting transplant surgeon and stored in a cardioplegic solution and transported to the University of Notre Dame with a transport/storage time of 1-8 hours. Donor hearts were then cut into region-specific pieces. Left ventricles were sectioned and decellularized as shown in Figure 1 A and B. Cell-free ECM is composed of the structural proteins as well as the secreted products of the resident cells [35]. Preservation of the ECM biochemical composition is crucial since ECM provides bioactive cues that affect cell response to its environment such as proliferation, migration, and differentiation [21,36,37]. Therefore, to ensure human cardiac tissue decellularization with the preservation of ECM bioactivity, we optimized previous decellularization methods [38,39] and used a combination of ionic and non-ionic detergent washes. Due to the fatty nature of the human heart, we also added an alcohol-based delipidation step following decellularization. By conducting hematoxylin and eosin (H&E) staining the absence of DNA was confirmed and Masson’s trichrome was used to show preservation of ECM proteins (Figure 1C). In addition, the double-stranded DNA content was measured and verified to be less than 50 ng/mg before using dhECM as bioink (Figure 1D). dhECM was then solubilized and mixed with the other hydrogels prior to printing.

**Figure 1.**
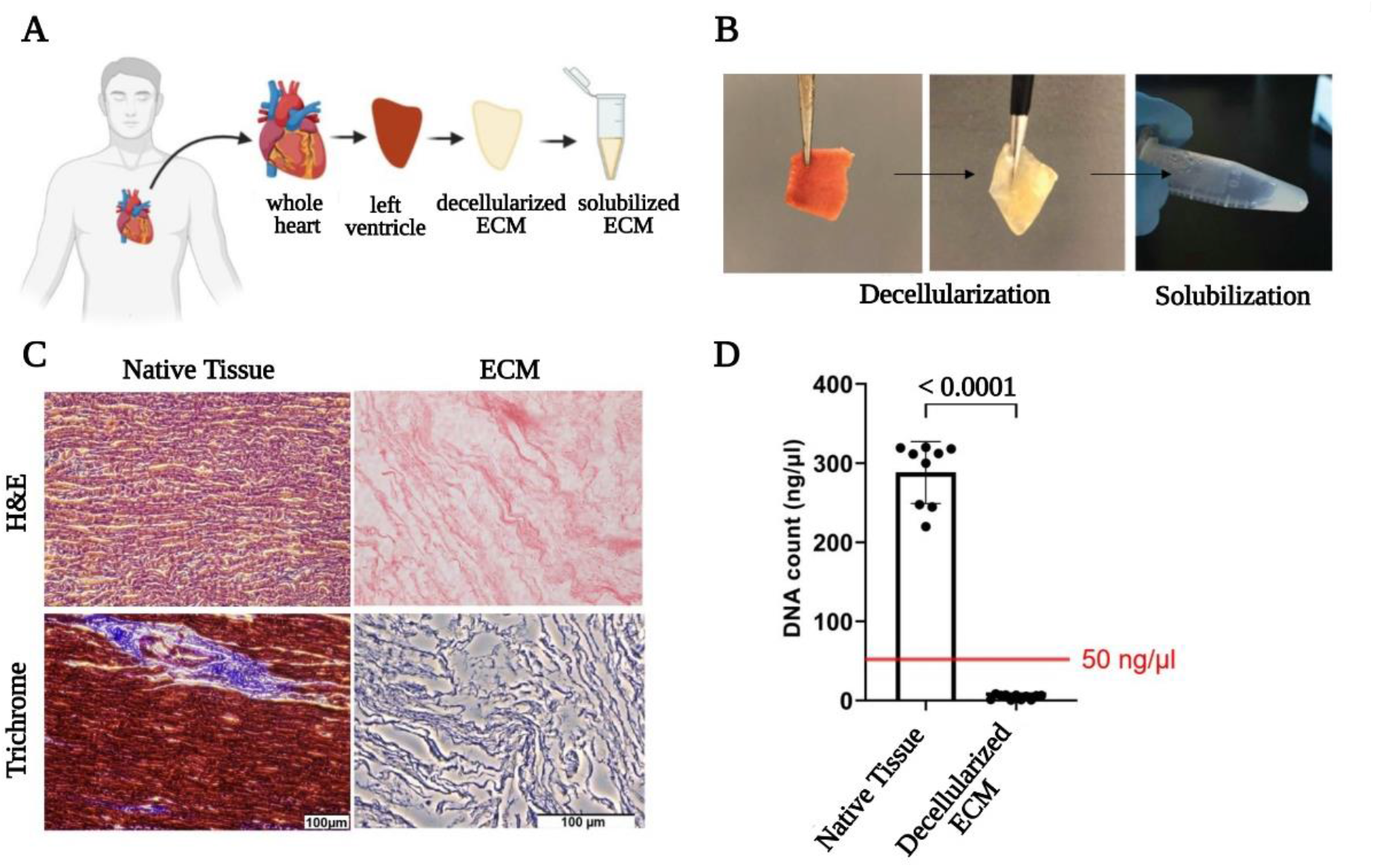
Decellularization of the native tissues and their biochemical analysis. **(A)** Schematic and **(B)** representative images demonstrating the workflow of the decellularization and solubilization process, **(C)** Hematoxylin and eosin (H&E) and Masson’s Trichrome staining of native tissue and ECM, **(D)** Quantitative measurements of total DNA content of native, decellularized and delipidized cardiac tissue.

### 2.2. Mechanical Characterization and Swelling

dhECM mixed with either GelMA alone or GelMA-MeHA yielded different material properties. Mixing dhECM with both polymers increased the Young’s Modulus significantly. Moreover, by including a second crosslinking step with mTGase, the mechanical properties of the hydrogel blends were further improved. Using a nanoindenter a uniaxial compression test was performed, and Young’s modulus of each hydrogel was determined (Figure 2A). The Young’s modulus of the hydrogels without and with mTGase treatment were measured to be 2.4 ± 0.4 kPa and 3.5 ± 0.5 kPa for G, 0.6 ± 0.1 kPa and 2.8 ± 0.7 kPa for GE, 14.4 ± 1.8 kPa and 24.5 ± 2.9 kPa for GM, 9.9 ± 2.6 kPa and 18.4 ± 2.8 kPa for GME, respectively (n≥3 for all). The improved elastic modulus of GM hydrogels is most likely resulting from the increased polymer concentration and crosslinking sites [40]. These results also demonstrate that mTGase treatment improved the Young’s modulus of the hydrogels which is in agreement with results shown in previous studies [32,41]. Moreover, constructing composite hydrogels by mixing dhECM with G and GM, allowed us to tune the Young’s modulus values in a wide range from 0.6 kPa to 18.4 kPa. This range can be further widened by changing the concentrations of GelMA and MeHA individually and mixing them in different ratios as shown previously [34].

**Figure 2.**
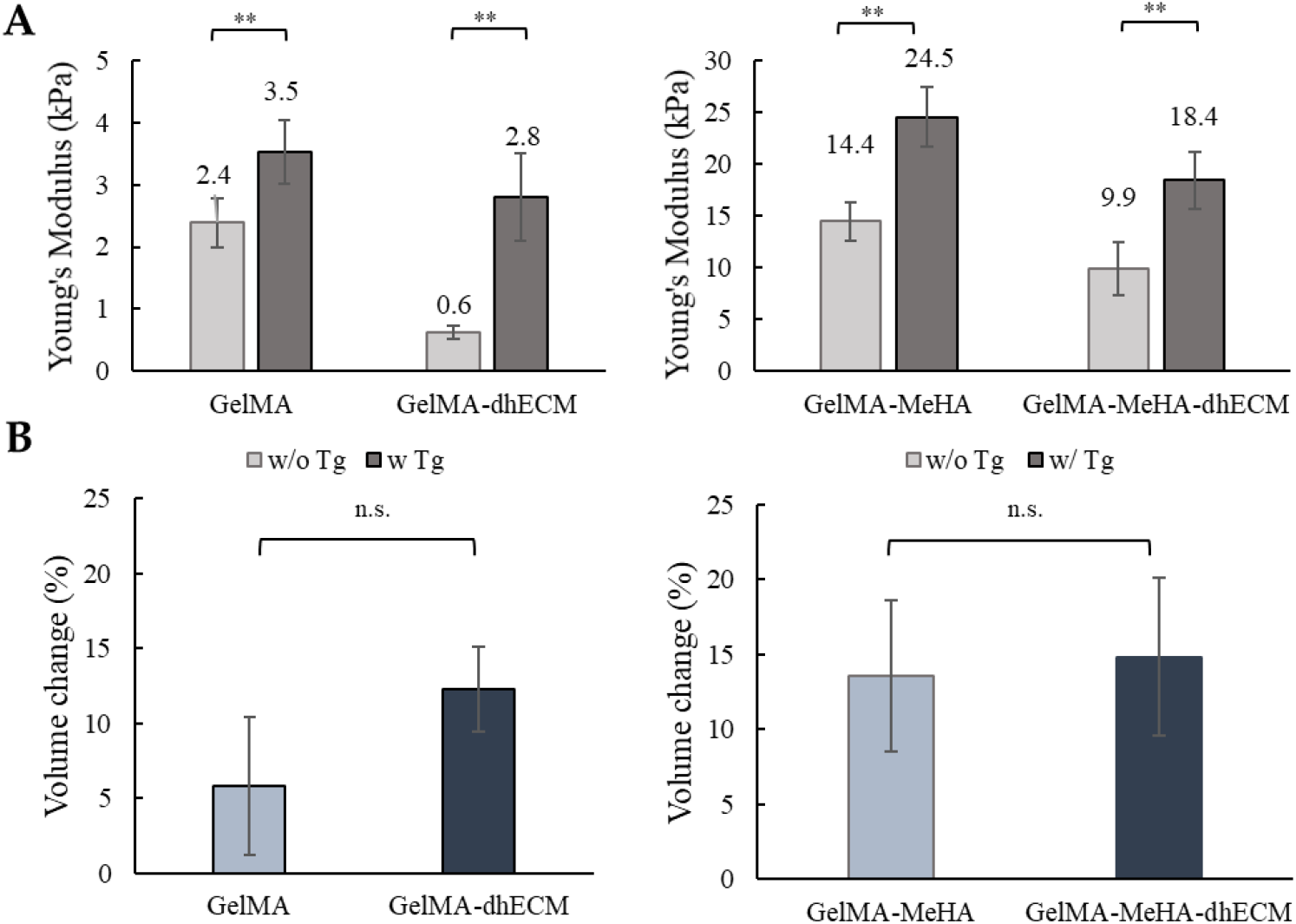
Mechanical and swelling characteristics of the hydrogels (**A**) Young’s modulus values of GelMA and GelMA-dhECM hydrogels without and with microbial Transglutaminase (mTGase) treatment (left), of GelMA-MeHA and GelMA-MeHA-dhECM hydrogels without and with mTGase treatment (right) (**B**) Swelling properties of GelMA and GelMA-dhECM hydrogels with mTGase treatment (left), of GelMA-MeHA and GelMA-MeHA-dhECM hydrogels with mTGase treatment (right). (** p < 0.01, n.s.: not significant) (Student’s t-test) (n≥3)

For the rest of this study, the two-step crosslinking method was used to create tissue constructs at a stiffness comparative to native human heart tissue (~10 kPa) [42]. For the swelling analysis, the volume of the hydrogels was measured after 24 hours incubation at 37 °C, allowing the gels to reach equilibrium swelling. G was observed to be swollen the least with 6% ± 4%, and the others show comparable swelling properties with 12% ± 3% for GE, 14% ± 5% for GM and 15% ± 5% for GME respectively as shown in Figure 2B. Previous studies have shown that introducing MeHA to low concentration GelMA hydrogels (<10%) lowers the swelling ratio, whereas remains ineffective in gels with greater than 10% GelMA, when only UV crosslinking was applied [34]. In this study, dual crosslinking allowed further crosslinking of GelMA without altering MeHA polymers. It is well-known that higher crosslinking density results in smaller pore sizes [40]. We hypothesize that this is the reason for the reduced swelling ratio in the GelMA gels and plan to expand upon these findings in future studies.

### 2.3. Rheological Characterization

In order to better ascertain the potential printability of the hydrogel blends, we utilized rheological experiments to measure various viscoelastic properties. First, the storage and loss modulus of each hydrogel blend was determined at a fixed frequency of 1 Hz and strain of 3%. GM hydrogel had the greatest storage and loss modulus with 8086.7 ± 83.8 Pa and 3203.1 ± 73.8 Pa, respectively. The storage and loss modulus of G and GE were nearly identical with storage modulus values of: 4895.7 ± 170.3 Pa, 4714 ± 16.8 Pa and with loss modulus values of: 2309 ± 34.2 Pa, 2304 ± 17.3 Pa respectively. Finally, GME had a storage modulus of 537.8 ± 14.0 Pa and a loss modulus of 295.6 ± 5.9 Pa (Figure 3A). To observe the relationship between storage and loss modulus with respect to frequency, the hydrogels were kept at a strain of 2% over a frequency range from 0.1 to 10 Hz. A slight decrease in the storage modulus and a slight increase in the loss modulus were observed in each group (Figure 3B). The storage and loss modulus of G and GM gels remained the same, indicating preserved solid-like structures. However, when hdECM was added to the GM, the decrease of the storage modulus and the increase of the loss modulus were more distinct, suggesting the increasing frequency weakened the link between the polymer chains. Finally, a flow ramp experiment was conducted for a shear range between 0.1 to 90 1/s and the shear rate-stress graphs were plotted, and viscosity was calculated from the slope. G and GE had similar viscosities whereas GM was significantly more viscous compared to GME (Figure 3C). As expected, viscosity for each hydrogel dropped with increasing shear rate.

**Figure 3.**
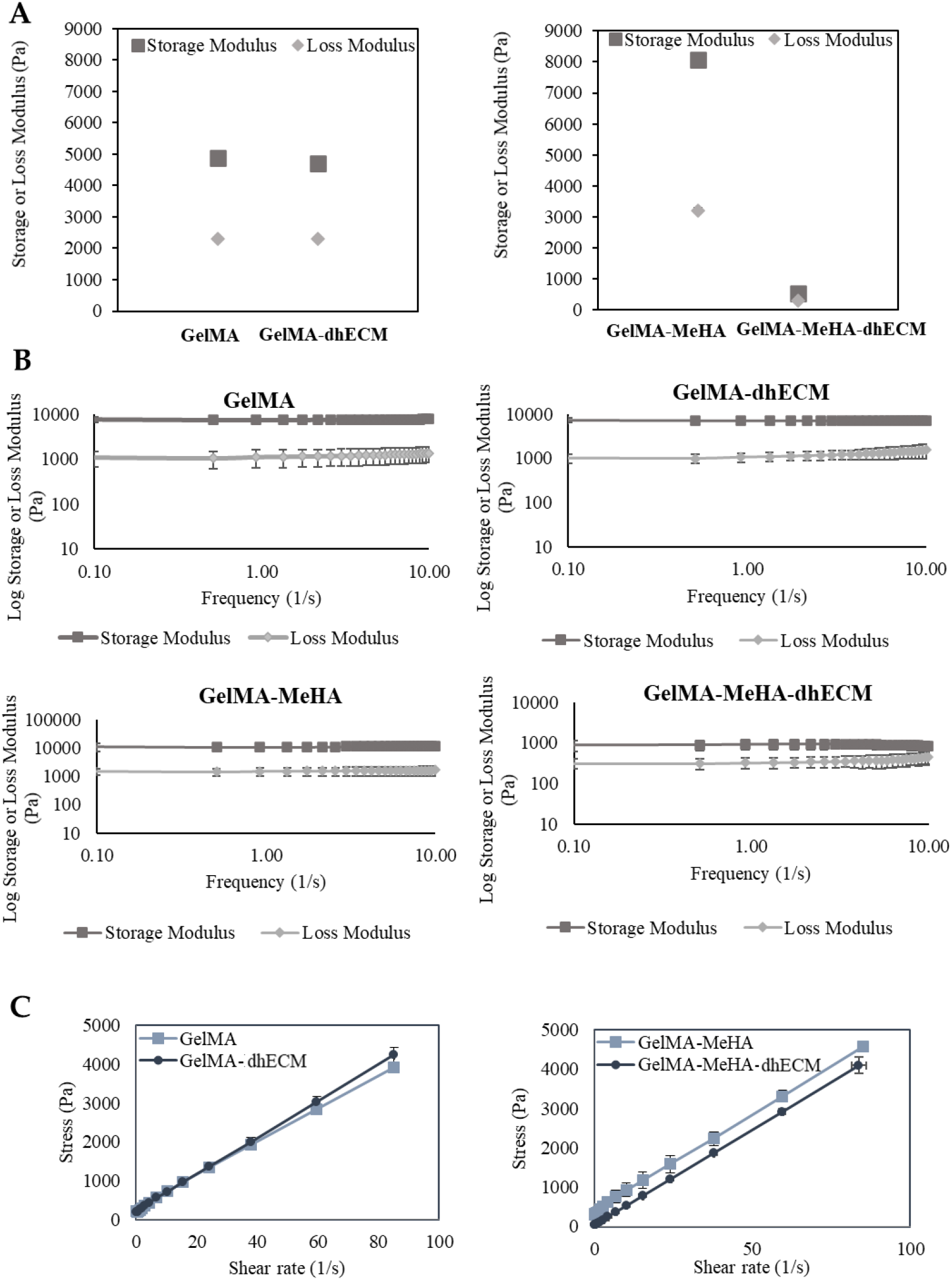
Viscoelastic properties of GelMA, GelMA-dhECM, GelMA-MeHA, GelMA-MeHA-dhECM hydrogels (**A**) Storage and lost modulus values at a constant frequency of 1Hz and a strain of 3%, (**B**) Change of storage and loss modulus with respect to frequency at a strain of 2%, (**C**) Stress-strain relationship revealing viscosity. (n≥3)

In the light of elastic modulus results, it was expected that GM and GME hydrogels would have superior viscoelastic properties compared to G and GE hydrogels. However, the rheology experiments revealed that GME hydrogels have the smallest storage modulus and viscosity. This could be explained by the fact that the Young’s modulus of polymer blends is less sensitive to the interaction and morphological changes than the yielding properties [43]. SEM images were taken to observe the microscopic structure of the hydrogels and GME constructs were distinctly more wrinkled than the other hydrogels (Figure S1).

### 2.5. 3D printing

To investigate the printability of the different hydrogels, 2-layered grid-shaped constructs (6 mm x 6 mm) were printed, with a 2 mm distance between neighboring lines. All hydrogel blends were printable, and the printed samples were dual crosslinked (Figure 4). The best printability was achieved with G which best preserved the distance between neighboring lines (Figure 4A). Using GE and GM, the grid-shaped construct was successfully printed as well (Figure 4B and C). It was observed that GME bioink required a longer time in the ice bath (~1.5-2 min) to reach the same consistency as the other bioinks (~1-1.5 min) prior to printing and its stability during printing was lower compared to the other blends which is in line with the rheology results (Figure 4D). A trend where the middle bottom square is greater than the middle top square can be seen for all materials. This might be due to the slight movement of the printing platform due to the vibration of the printhead. The effect of this vibration was minimized by placing parafilm both on the stage and inside the dish under the glass (Figure S2). Previous studies reported the printability of decellularized cardiac dECM alone [22], or dECM composites with PEGDA [4], HA, gelatin and various crosslinkers [13], only MeHA [33] and only GelMA [44]. By mixing GelMA with dhECM and MeHA, our group was able to develop a novel, composite hydrogel that resulted in better printability compared to the printability of dECM [22] and MeHA [33] alone. Moreover, the pre-designed shape can be printed more accurately if a higher gauge nozzle was used [4]. Additionally, printing speed and pressure can be adjusted to improve the printability; yet, these parameters affect the cell viability [45]. In our study, we investigated the cell viability when they were printed using a 22G nozzle with a translational speed of 75 mm/min and with a printing pressure of 20-30 kPa. The individual effect of these parameters on cell viability can be further investigated in future studies.

**Figure 4.**
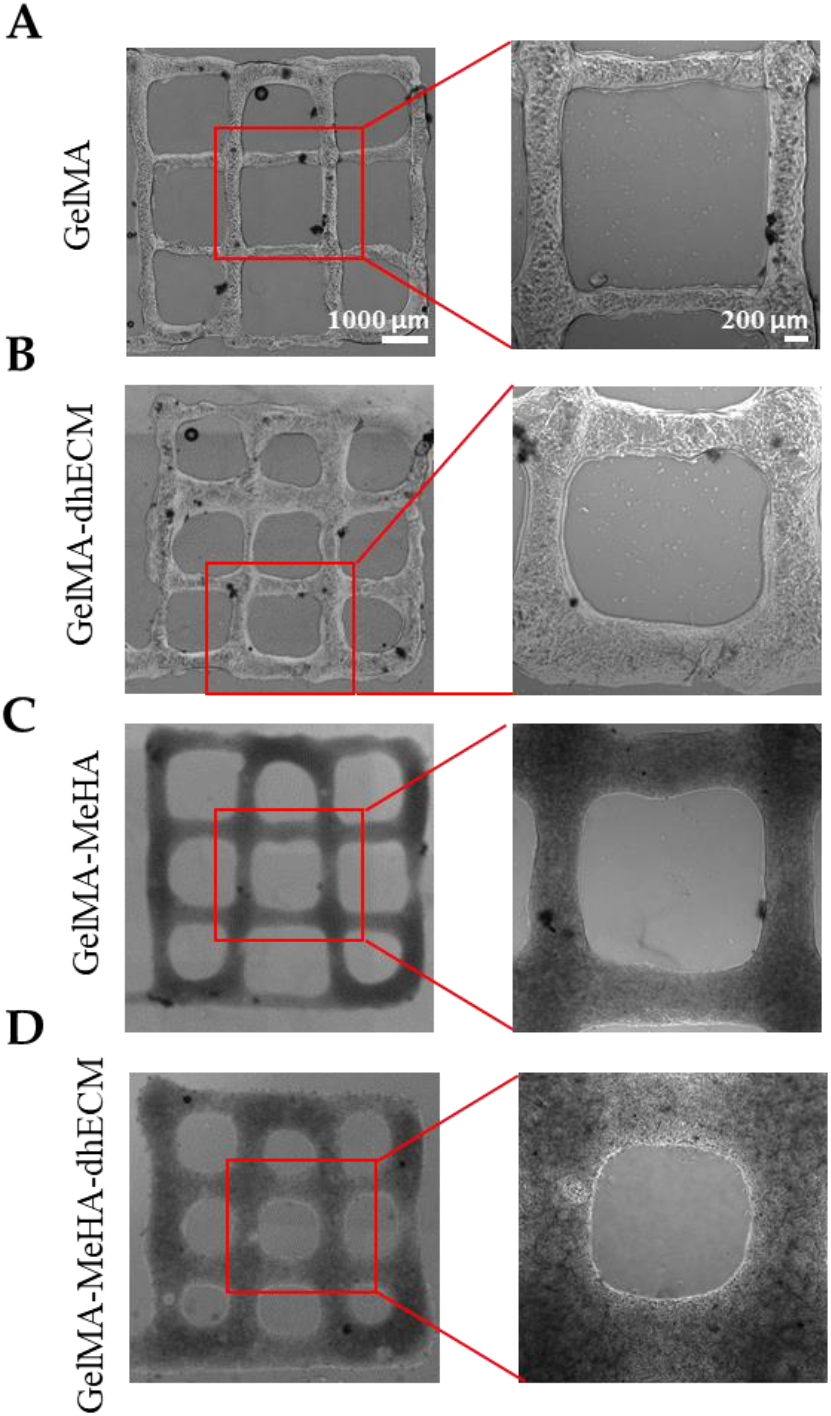
Brightfield images of 3D printed 2 layered-grid constructs using (**A**) GelMA, (**B**) GelMA-dhECM, (**C**) GelMA-MeHA, (**D**) GelMA-MeHA-dhECM hydrogels. (n≥3)

### 2.6. Cell viability

To evaluate the effect of the bioink composition on cell viability, iCM and hCFs were printed with G and GM with or without dhECM supplementation. To recapitulate the healthy cardiac tissue, iCMs were printed with low stiffness G or GE bioinks (3.5 or 2.8 kPa) whereas hCFs were printed with high stiffness GM or GME bioinks (24.5 or 18.4 kPa) to represent the scar tissue. The day after printing, more than 65% viability was achieved for each hydrogel (Figure 5). The percentage of live cells was calculated as 67% ± 6% and 74% ± 10% for iCMs printed in G only or GE bioinks, respectively (Figure 5 A, B and E). The viability of hCFs was 79% ± 3% in GM and 84% ± 3% when the cells were mixed with GME bioink prior to printing (Figure 5 C, D, and F). Combining dhECM with G slightly improved iCM viability and a similar effect was observed for hCFs when dhECM was combined with GM hydrogel.

**Figure 5.**
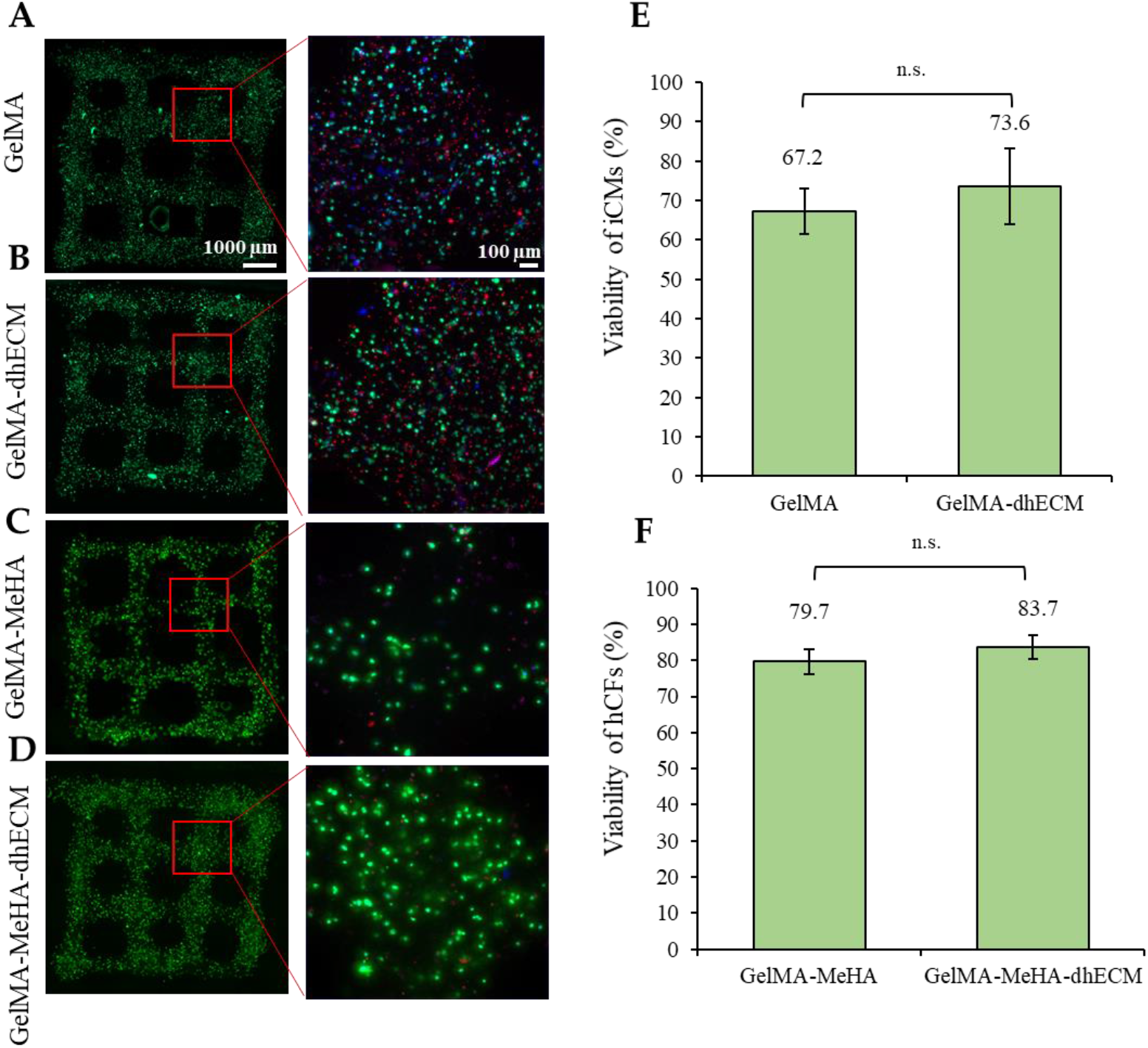
Cell viability analysis of printed cell-laden hydrogels prepared using (**A**) GelMA, (**B**) GelMA-dhECM, (**C**) GelMA-MeHA, (**D**) GelMA-MeHA-dhECM, (**E**) live cell percentage comparison for GelMA and GelMA-dhECM (**F**) live cell percentage comparison GelMA-MeHA and GelMA-MeHA-dhECM. (Student’s t-test) (n≥3)

The iCM viability reported in this study was slightly lower compared to previous studies in which the cells were printed using an extrusion-based printer with different bioinks, which may be resulting from factors such as UV light treatment, mixing the cells with a photoinitiator (PI), and encapsulated cell density. As Koti et al. reported in their study, UV and PI are factors that are negatively affecting the cell viability along with the increasing gel concentration [12]. However, to meet the native tissue like mechanical properties using G and GM hydrogels, the addition of UV-sensitive PI and UV treatment are necessary while the hydrogels obtained using visible light crosslinking were measured to have Young’s modulus of less than 1kPa (Figure S3). Lee et al. printed highly concentrated iCMs and collagen in different printheads, using collagen as supporting material in a gel bath and reported the viability only immediately after printing as over 95% [14]. The difference observed in viability might be resulted from printing in a gel bath, not mixing the cells with any bioink or PI, high printing cell concentration, and assessing the viability immediately after printing. In another study, Kumar et al. used a fibrinogen-gelatin based hydrogel with a visible light induced PI to print iCMs and reported cell viability over 90% [16]. The difference in viability could be resulting from avoiding employing the UV-induced PI and polymerization with UV-light in their process. Despite the ~70% viability quantified in this study, healthy tissue-like beating of the cells was achieved (Movie S1 (iCMs printed with G) and S2 (iCMs printed with GE)). Better viability and beating of the cells was observed when they were in close proximity with high cell-to-cell interaction, and when they were separately encapsulated the chance of survival decreased, in agreement with our previous study [46]. The viability of the encapsulated iCMs might be improved by optimizing the cell density; however, this will disturb the homogeneity of the cell-gel mixture, affecting the printing ability and quality negatively. One other solution to improve viability might be introducing a perfusion system allowing the media to reach through the thick tissue [14]. These and other methods could be explored to increase viability in future studies.

### 2.7 Beating and Phenotypical Characterization of Printed Constructs

Spontaneous beating was observed as soon as 24 hours after printing. To investigate the beating characteristics of iCMs, brightfield videos were taken 9 days after printing (Movie S3 (iCMs printed with G) and S4 (iCMs printed with GE)). Beating velocity and frequency of the printed tissue constructs were quantified by video analysis of lateral displacement of spontaneous beating iCMs and heat maps were generated using a custom made MatLab code (Figure 6A and B). Beating velocity was measured to be 7.7 μm/s ± 3 μm/s for G and 8.0 μm/s ± 2 μm/s for GE (Figure 6C) and beating frequency was calculated as 0.54 Hz ± 0.30 Hz for G and 0.65 Hz ± 0.26 Hz for GE (Figure 6D). As the results indicate, dhECM slightly improved beating kinetics and yielded faster iCM beating at higher frequencies. In previous studies, the beating velocities of the printed cardiac constructs were not reported. However our results are in perfect alignment when iCMs were directly encapsulated in collagen hydrogel and the beating velocity was reported as 8 μm/s, and frequency as 0.67 Hz [47].

**Figure 6.**
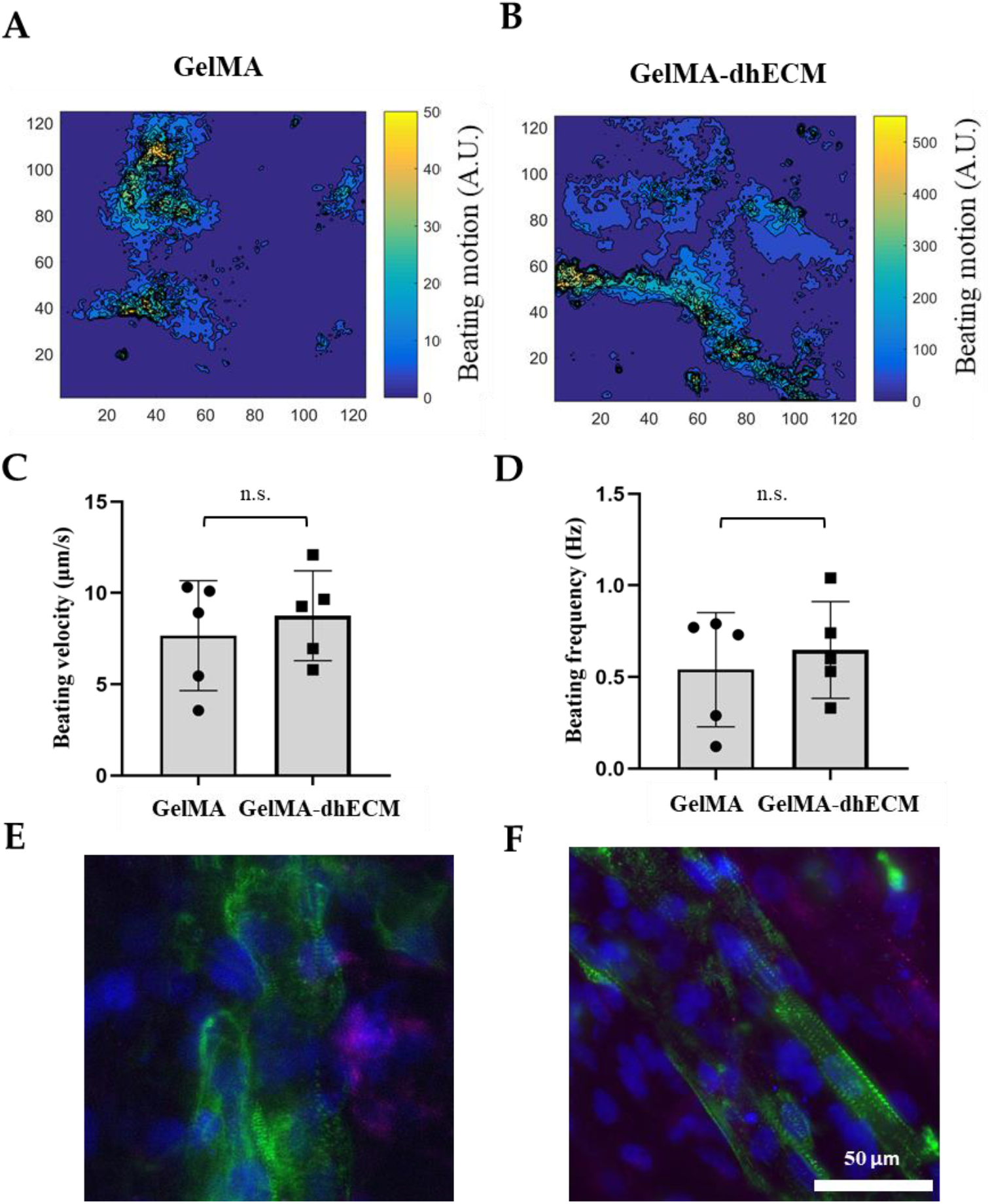
Beating and phenotypical characterization of iCMs. The heat maps showing the beating velocity magnitude (A.U.) and distribution of iCM (**A**) when they were printed in GelMA hydrogel, (**B**) when they were printed in GelMA-dhECM hydrogel, (**C**) average beating velocity of iCMs, (**D**) average beating frequency of iCMs. Immunostaining of iCMs for CX43 (magenta) and sarcomeric alpha-actinin (green), (**E**) when they were printed in GelMA hydrogel, and (**F**) when they were printed in GelMA-dhECM hydrogel. Cell nuclei are stained with DAPI (blue). (Student’s t-test) (n≥3)

On day 21 after printing, Connexin 43 (CX43), a canonical cardiac intercellular junction protein, and sarcomeric alpha-actinin, a cytoskeletal actin-binding protein, expression were investigated by immunostaining. For both G and GE hydrogels, iCMs showed CX43 expression and striated sarcomeric alpha-actinin structure (Figure 6 E and F). However, in the presence of hdECM, cells were observed to be more physiological with more elongated and organized striations (Figure 7F). Similar organized striation of sarcomeric-alpha-actinin was observed in previous studies when the iCMs were patterned like native myocardium, which is important for developing higher contractile force, and also an indication of more mature cells [48,49].

**Figure 7.**
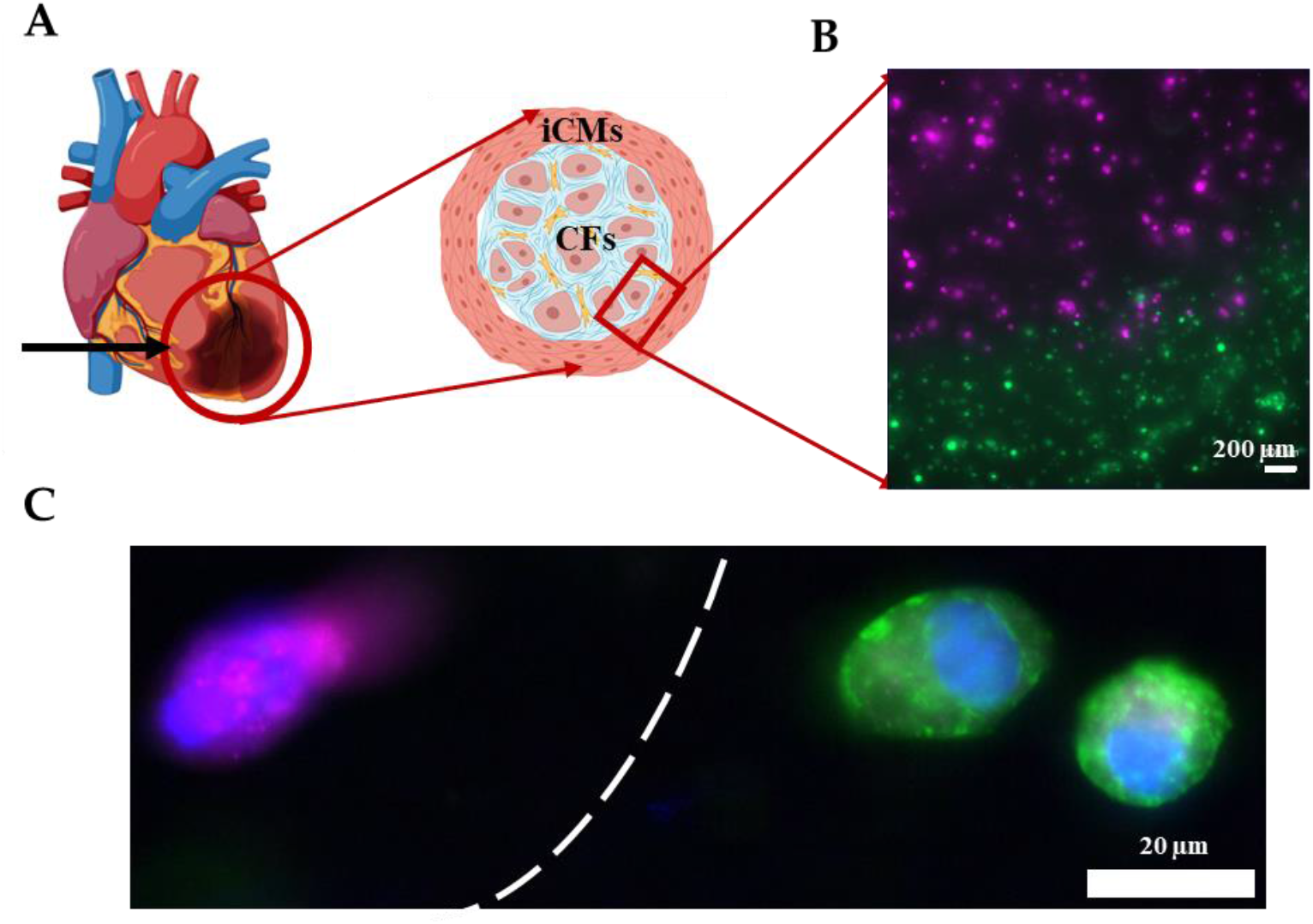
Printing the infarct region (**A**) Schematic showing the infarct region model and corresponding cells (**B**) Image showing the successful printing of the infarct region with iCMs (green) and hCFs (magenta), (**C**) Immunostaining of iCMs for sarcomeric alpha-actinin (green) and hCFs for Vimentin (violet). Cell nuclei are stained with DAPI (blue).

### 2.8. 3D printing the infarct boundary region

As shown in the previous section, the Young’s modulus of GM and GME hydrogels were approximately an order of magnitude different from each other. This tunability in stiffness allows these materials to be used in a variety of applications. It is known that post-MI ECM remodeling yields scar tissue *in vivo* [50]. This tissue is significantly stiffer compared to healthy cardiac tissue and consists mostly of CFs and myofibroblasts [51]. As a proof of concept, an infarct boundary region was printed using GME mixed with hCFs to model the infarct region surrounded by the healthy tissue which is modeled using GE bioink mixed with iCMs as shown in Figure 7A. This model has been used to demonstrate the infarct boundary region in previous studies [3,52]. Prior to printing, iCMs were tagged with a green cell tracker and hCFs were tagged with a deep red cell tracker (Figure S4). Fluorescent images were taken immediately following printing, showing the printing success with dual printheads, different materials and cell types (Figure 7B). Immunostaining was performed on the printed products to demonstrate that the CFs were expressing fibroblast protein Vimentin and iCMs were expressing cardiac protein sarcomeric alpha-actinin (Figure 7C). Like our study, Koti et al. also printed using dual printhead to print “spatially defined 3D constructs” using rat CMs and CFs. Our study advances the field by introducing printing by using hydrogels supplemented with dhECM and human-derived cardiac cells and achieving various mechanical properties to better mimic the infarct region. Although the model achieves the order of magnitude difference in stiffness of healthy tissue and scar tissue, it lacks achieving the physiologically accurate stiffness of myocardial tissue which remains as a limitation and will be addressed in future studies.

## 3. Conclusions

In this study, we showed the extrusion based 3D printing of GelMA and GelMA-MeHA with the addition of human cardiac tissue ECM. We have fully characterized the mechanical properties of these bioinks and evaluated cell compatibilities. We reported that the elastic modulus of GME hydrogel was an order of magnitude greater compared to GE hydrogel. Leveraging this stiffness difference, we demonstrated as a proof of concept the potential use of GE and GME hydrogels with iCMs and hCFs, respectively, in mimicking post-MI cardiac tissue.

The fabricated 3D *in vitro* MI models can be used to study MI recovery and potential therapies including cardiac patches and cell injection with human cells. This model can also be used to better understand the relation between scar tissue and healthy tissue. Moreover, by mixing the iCMs and hCFs in different concentrations, better modeling of the infarct boundary region can be achieved. These newly developed bioinks and printing method can potentially provide researchers with tunable, printable hydrogels that can be used to mimic human physiological conditions *in vitro*, providing a crucial bridge between 2D *in vitro* and expensive long *in vivo* and clinical studies.

## 4. Materials and Methods

### 4.1. Ethics statement

De-identified human hearts that were deemed unsuitable for transplantation and donated to research, were acquired from Indiana Donor Network under the Institutional Review Board (IRB) approval for deceased donor tissue recovery. All human tissue collection conformed to the Declaration of Helsinki.

### 4.2. Decellularization and Solubilization of Human Cardiac Tissue

Hearts were cut into region specific pieces and stored at −80 °C until further use. Left ventricles from donors were sectioned and decellularized using 1% Sodium dodecyl sulfate (SDS, VWR, PA, USA) for 24 h or until the tissues turned transparent white. Then, samples were transferred to 1% Triton X-100 (Sigma-Aldrich, MO, USA) for 30 min. After decellularization, samples were washed thoroughly with deionized (DI) water to remove any residual detergent. To delipidize, dECMs were washed in isopropanol for 1 hour and rehydrated via DI washes. All steps were conducted with constant agitation at room temperature (RT).

dECMs were digested in a 1 mg/mL pepsin in 0.1 M HCl at RT with constant stirring until a homogeneous solution was obtained. The insoluble remnants were removed by centrifugation, the supernatant was neutralized using 1 M NaOH solution, and either used immediately or flash frozen and stored at −80°C until use. Prior to experiments, the total protein concentrations was measured using Rapid Gold BCA Assay (Thermo Scientific, MA, USA) and dECM solutions were diluted to 3 mg/mL with Phosphate-Buffered Saline (PBS, Corning, NY, USA).

### 4.3. GelMA and MeHA synthesis

GelMA synthesis was performed by following a previously established protocol [53]. Briefly, 10 g of gelatin (gel strength 300 g Bloom, Type A, from porcine skin, Sigma-Aldrich, St. Louis, MO, USA) was dissolved in 100 mL of phosphate-buffered saline (PBS, Corning, NY, USA) at 60 °C. 2 mL of methacrylic anhydride (MAnh, Sigma-Aldrich, St. Louis, MO, USA) was added dropwise and the pH was adjusted to 8. The solution was kept at 60 °C for 3 hours with constant stirring. Then the solution was diluted with 400 mL of PBS (pre-warmed to 40-50 °C) and stirred for 15 mins. The solution was then transferred into 12-14 kDa MWCO dialysis membranes (VWR, Chicago, IL, USA) and dialyzed against DI water for one week, with twice daily water changes, before filtering and lyophilizing.

For MeHA synthesis, 0.2 g of HA (MW: 1.5-1.8 x 10^6^ Da, Sigma-Aldrich, St. Louis, MO, USA) was dissolved in 60 mL DI water at room temperature with constant stirring overnight. The next day 40 mL dimethyl formamide (DMF, Sigma-Aldrich, St. Louis, MO, USA) was added dropwise using a glass pipette. Then, 0.8 mL of MAnh was added dropwise and the pH was set to 8-9 for the reaction to take place at 4 °C overnight. On the next day, the mixture was placed in 12-14 kDa MWCO dialysis membranes and dialyzed against DI water for 3 days, by replacing the water 2-3 times a day. Finally, the MeHA solutions were filtered and lyophilized for further use.

### 4.4. Cell Culture

#### 4.4.1. hiPSC Culture and iCM differentiation

DiPS 1016 SevA hiPSC line derived from human skin fibroblasts were seeded and kept in culture on Geltrex (1% Invitrogen, USA)-coated culture flasks using mTeSR (StemCell Technologies, Vancouver, BC, Canada) supplemented with 1% penicillin (Pen) (VWR, USA) at 37 °C with 5% CO^2^. When confluency of 80% was reached, the hiPSCs were detached using Accutase (StemCell Technologies, Vancouver, BC, Canada), and seeded in culture well plates in mTeSR1 media supplemented with Rho-associated, coiled-coil containing protein kinase (ROCK) inhibitor (5 μM, StemCell Technologies, Vancouver, BC, Canada). Until 95% confluency was reached, the culture was maintained with daily media changes.

A previously established protocol was adapted to differentiate iCMs from hiPSCs [51]. Briefly, when the hiPSCs reached 95% confluency, they were treated with RPMI Medium 1640 (Life Technologies, USA) supplemented with B27 without insulin (2%, Invitrogen, USA), beta-mercaptoethanol (final concentration of 0.1 mM, Promega, USA) and Pen (1%) (CM (-)) with the addition of Wnt activator, CHIR99021 (CHIR) (12 μM, Stemgent, USA). Exactly twenty-four hours later, media were replaced with CM (-) without CHIR. On day 4, iCMs were treated with CM (-) media supplemented with Wnt inhibitor IWP-4 (5 μM, Stemgent, USA). Media was changed back to CM (-) on day 6. Three days later (day 9), media was replaced with RPMI Medium 1640 supplemented with B27 (2%, Invitrogen, USA), beta-mercaptoethanol (final concentration of 0.1 mM), and Pen (1%) (CM (+)). After day 9, media was changed every 3 days, and beating was observed generally by day 21 of differentiation as stated in previous papers [52–54].

#### 4.4.2. Human CF culture

Human cardiac fibroblasts (Cell Applications, San Diego, CA, USA) were kept in culture using DMEM High Glucose (Corning, Corning, NY, USA) supplemented with fetal bovine serum (FBS, 10%) (Hyclone, South Logan, UT, USA), penicillin-streptomycin (P/S, 1%) (Gibco, Waltham, MA, USA) and SD208 (3μM, TGF-β pathway inhibitor) (Sigma-Aldrich, St. Louis, MO, USA) at 37 °C with 5% CO^2^. The culture was maintained with daily half media changes until they reached approximately 80% confluency.

### 4.5. Preparation of Hydrogels for Material Characterization

Four different hydrogel solutions were prepared: G, GE, GM, GME. G solution was prepared by dissolving G (10% *w/v*) and adding Irgacure 2959 PI (0.05% *w/v*, Sigma-Aldrich, St. Louis, MO, USA) in PBS. The solution was kept at 37 °C until G was completely dissolved. To prepare the GE solution, G was prepared as explained above and mixed thoroughly with dhECM to achieve a final concentration of 10% *w/v* of G and 1 mg/mL of dhECM. MeHA was dissolved in PBS (2% *w/v*), at 80 °C for 1 hour. Then it was mixed with G (20% *w/v*) and PI (0.2% *w/v*). To achieve a better mixture of GM hydrogels, the mixture was placed in an Eppendorf tube with a stir bar on a stirrer at 37 °C for an hour. Finally, GME solution was prepared similarly, and dhECM was added after the GelMA and MeHA mixed completely and final concentrations of 1mg/mL, 10% *w/v*, 1% *w/v*, and 0.1% *w/v* was achieved for dhECM, GelMA, MeHA and PI, respectively. The microbial transglutaminase (mTGase, Modernist Pantry, Portsmouth, NH, USA) solution was prepared in PBS (80 mg/mL *w/v*) and kept at 37 °C until mTGase was completely dissolved.

After all the solutions were prepared, the hydrogels were prepared by transferring 100 μl from each solution onto a stage, in-between 1 mm thick spacers, and a glass slide was placed on top of the solution to achieve the required thickness. The gels were then exposed to 6.9 mW/cm^2^ UV irradiation by using a UV lamp (Lumen Dynamics, (Mississauga, ON, Canada). The gels were treated with mTGase solution for 30 mins at 37 °C, then the mTGase solution was replaced with PBS. The gels were kept at 37 °C overnight to achieve equilibrium swelling of the hydrogels.

### 4.6. Materials Characterization

#### 4.6.1. Mechanical Properties

To measure the Young’s modulus of each hydrogel, a compression test was conducted using a nanoindenter (Optics 11, Westwood, MA, USA) with an indentation probe (spring constant of 0.51 N/m, tip diameter of 46 μm) as described previously [54]. Young’s modulus was calculated as the slope of the stress-strain curve in the elastic region, which was determined using an in-house MATLAB code.

#### 4.6.2. Swelling Test

By measuring the volume change between the freshly prepared hydrogels and those that were incubated in PBS solution at 37 °C for 24 hours, the swelling properties of the hydrogels were determined. To calculate the volume of the hydrogels, the surface area was multiplied by the thickness. To measure the surface area of the hydrogels, images of the gels were taken immediately after preparation and 24 hours later and analyzed using ImageJ (National Institutes of Health Bethesda, MD, USA). The gels were prepared to have an initial thickness of 1 mm. The next day, the thickness of the gels were determined using the force reading in the rheometer, which showed zero up until the upper plate touched the surface of the gel and started increasing after contact occurred. The thickness reading at this contact point was used as the thickness of the gel.

#### 4.6.3. Rheological Properties

Viscoelastic properties of the hydrogels were characterized using an HR-2 Hybrid Rheometer (TA Instruments, New Castle, DE, USA) with 8 mm diameter parallel-plate geometry. First, the storage (G’) and loss modulus (G’’) of the hydrogels were recorded for 1 min at 37 °C, under a fixed frequency of 1 Hz and strain of 3%. Then, the storage and loss modulus of the hydrogels were recorded at 2% strain over a frequency range from 0.1 to 10 Hz at 37 °C, to observe the viscoelastic properties of the hydrogels. Finally, shear stress and viscosity of each hydrogel were recorded for a shear rate change from 0 to 90 1/s.

#### 4.6.4. Printability

The hydrogels were prepared as previously described. After preparation, each hydrogel was transferred into a cartridge and placed in an ice bath for 1-2 mins to achieve the required consistency for printing. A CELLINK Inkredible+ Bioprinter (CELLINK, Gothenburg, Sweeden) was used for printing. A grid pattern consisting of two layers with dimensions 0.6 cm x 0.6 cm was printed on a charged glass placed in a 35 cm dish (Figure S2), using a 22G nozzle with a translational speed of 75 mm/min. Right after printing, the constructs were treated with 30 s UV (6.9 W/cm^2^ UV radiation) using a UV lamp (Lumen Dynamics, Mississauga, ON, Canada). Printed constructs were treated with mTGase solution for 30 mins and kept at 37 °C overnight. The next day brightfield images were taken.

#### 4.7. Printing the tissue constructs

##### 4.7.1. Viability assessment

iCMs were reseeded on culture day 30-50. On day 3 the cells were collected using Trypsin ethylenediamine tetraacetic acid (EDTA) (VWR, Chicago, IL, USA). Cells were mixed with either GelMA (10% *w/v* final) or GelMA (10% *w/v* final) – dhECM (1 mg/mL final) hydrogels with 0.05% PI to have a final density of 20 mil/mL.

hCFs were kept in culture until they reached 90% confluency in T75 flask. They were then collected using Trypsin EDTA. Cells were mixed with either GelMA (10% *w/v* final) – MeHA (1% *w/v* final) or GelMA (10% *w/v* final)-MeHA (1% *w/v* final) −dhECM (1 mg/mL final) hydrogels with 0.1% PI to have a final density of 1 mil/mL.

The samples were prepared as described in the previous section and placed in a cartridge (500 μL total volume). The same printing protocol was followed, and the printed constructs were washed with PBS. The constructs with iCMs were then washed with CM+, and the constructs with hCFs were washed using DMEM Complete media for 5 minutes, prior to mTGase (80 mg/mL *w/v* prepared in cell specific culture media) treatment for 30 mins. After 30 mins mTGase was removed and the constructs were kept in cell specific media for 15 mins before changing to fresh media and kept in culture.

The next day after printing, live/dead assay was performed (Life Technologies, Carlsbad, CA, USA) following the manufacturer’s instructions. Briefly, the constructs were washed with PBS and incubated at 37 °C for 30 mins in a solution containing Calcein AM (live cells, green, 2μM), Ethidium homodimer-1 (dead cells, red, 4 μM), and Hoescht (Thermo Scientific, MA, USA). For each construct, z-serial images were then taken with a fluorescence microscope (Zeiss, Hamamatsu ORCA flash 4.0, Thornwood, NY, USA). Live and dead cells were counted in ImageJ software. Live cell percentage was calculated by using Equation 3.

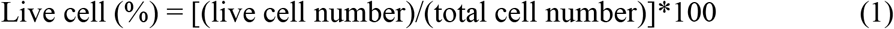

##### 4.7.2. Beating and Phenotypical Characterization

To analyze the contractility of iCMs, a block-matching algorithm was performed using MATLAB as described previously [42]. By using this method on the brightfield videos, the beating velocity and frequency of the iCMs were calculated and the heat maps were plotted.

For immunostaining, the 3D printed constructs were kept in culture for 3 weeks before fixing using paraformaldehyde (4%, Electron Microscopy Sciences, USA) for 45 minutes at RT. After fixing, they were washed with PBS and then permeabilized in Triton X-100 (0.1%, Sigma-Aldrich, USA) for 45 min. They were washed with PBS and blocked using goat serum (10%, Sigma-Aldrich, USA) for 2 hours. After blocking, constructs were incubated with sarcomeric alpha-actinin (ab9465, Abcam, United Kingdom), and Connexin 43 (CX43) (ab11370, Abcam, United Kingdom) primary antibodies diluted 1:200 and 1:100 respectively in goat serum at 4 °C overnight. The next day, the constructs were washed thoroughly with PBS and then incubated with Alexa Fluor 647 (A21245, Life Technologies, USA) and Alexa Fluor 488 (A11001, Life Technologies, USA) diluted 1:200 in goat serum at 4 °C for 6h. After incubation, constructs were washed with PBS until no background was seen. The samples were then fixed using ProLong Gold mounting medium (Thermo Scientific, MA, USA) with DAPI (ab 104139, Abcam, United Kingdom). Imaging was then performed using a fluorescence microscope (Zeiss, Hamamatsu ORCA flash 4.0, Thornwood, NY, USA).

##### 4.7.3. Printing the Infarct Boundary Zone

Bioinks were prepared by mixing iCMs in GE and hCFs in GME hydrogels as described above. Prior to mixing, the cells were tagged using Cell Tracker green (C2925) and deep red (C34656), (Life Technologies, Carlsbad, CA, USA) by following the manufacturer’s instructions. Using a custom-made G-Code the infarct region was printed using two printheads. The next day images were taken with a fluorescence microscope (Zeiss, Hamamatsu ORCA flash 4.0, Thornwood, NY, USA). For immunostaining, the printing was performed without mixing the cells with Cell Tracker. On the next day, immunostaining was performed on the printed infarct region constructs following the protocol given above (for Vimentin (ab137321, Abcam, United Kingdom, was diluted 1:200). Imaging was performed using a fluorescence microscope (Zeiss, Hamamatsu ORCA flash 4.0, Thornwood, NY, USA).

#### 4.8. Statistical analysis

For all replicates, the mean ± standard deviation (SD) was reported. To find any statistically significant differences, one-way analysis of variance (ANOVA) followed by Tukey’s post hoc was used. For comparing two individual groups, student’s t-test was used. All p-values reported were two-sided and p<0.05 was considered statistically significant. Sample size (n)≥3 for all individual experiments.

## Supplementary Materials

The following are available online at www.mdpi.com/xxx/s1, Figure S1: SEM images of the hydrogels, Figure S2: Parafilm coating (A) on the printing stage, (B) in the dish under the glass, Figure S3: Young’s Modulus of hydrogels with visible light crosslinking, Figure S4: Printed boundary region using iCMs in GelMA-dhECM bioink (green), and hCFs encapsulated in GelMA-MeHA-dhECM bioink (magenta), Video S1: Tissue like beating of iCMs encapsulated in GelMA. Video S2: Tissue like beating of iCMs encapsulated in GelMA-dhECM. Video S3: Brightfield video of beating of iCMs encapsulated in GelMA taken 9 days after printing. Video S4: Brightfield video of beating of iCMs encapsulated in GelMA-dhECM taken 9 days after printing.

## Author Contributions

Conceptualization, G.B and P.Z.; methodology, G.B, S.G.O. and P.Z.; software, G.B.; validation, G.B., S.G.O., B.W.E., and P.Z.; formal analysis, G.B.; investigation, G.B.; resources, P.Z.; data curation, G.B. and S.G.O.; writing—original draft preparation, G.B.; writing—review and editing, G.B., S.G.O., B.W.E., and P.Z.; visualization, G.B.; supervision, P.Z.; project administration, P.Z.; funding acquisition, P.Z. All authors have read and agreed to the published version of the manuscript.

## Funding

This research was funded by NIH Award #1 R01 HL141909-01A1, NSF Award #1651385, and the Naughton Fellowship.

## Acknowledgments

Figure 1A and Figure 7A are created with BioRender.com.

## Conflicts of Interest

The authors declare no conflict of interest.

## Notes

### Competing Interest Statement

The authors have declared no competing interest.

