## Supplementary material for "Tunable human myocardium derived decellularized extracellular matrix for 3D bioprinting and cardiac tissue engineering": Supplementary Materials.pdf

**This PDF includes:**
Materials and Methods
Figures S1 to S4

**Other supplementary materials for this manuscript includes:**
Movies S1 & S2

**Materials and Methods:**
**Hydrogel Preparation with Visible light Crosslinker**

Ruthenium (Ru) and Sodium Persulfate (SPS) solutions were prepared following the manufacturer's instructions (Advanced BioMatrix, Carlsbad, CA, USA). Simply, Ru was dissolved in 1X PBS at a concentration of 37.4 mg/mL and SPS was dissolved in 1X PBS at a concentration of 119 mg/mL. GelMA solution was prepared by dissolving GelMA (10% *w*/*v*) in PBS and Ru (2% *w*/*v*) and SPS (2% *w*/*v*) was included. Similarly, GelMA-MeHA solution was prepared by dissolving GelMA (20% *w*/*v*) and MeHA (2% *w*/*v*) in PBS and by mixing them in 1:1 ratio. Then Ru (2% *w*/*v*) and SPS (2% *w*/*v*) was included in the hydrogel mixture. 5 µL hydrogel solution was then placed on a stage in-between 100 µm thick spacers, and a glass slide was placed on top of the solution to achieve the required thickness. The hydrogel solution was then exposed to blue light for 3 s. Half of the gels were treated with mTGase solution for 30 mins at 37 °C, then the mTGase solution was replaced with PBS. The gels were kept at 37 °C overnight to achieve equilibrium swelling of the hydrogels.

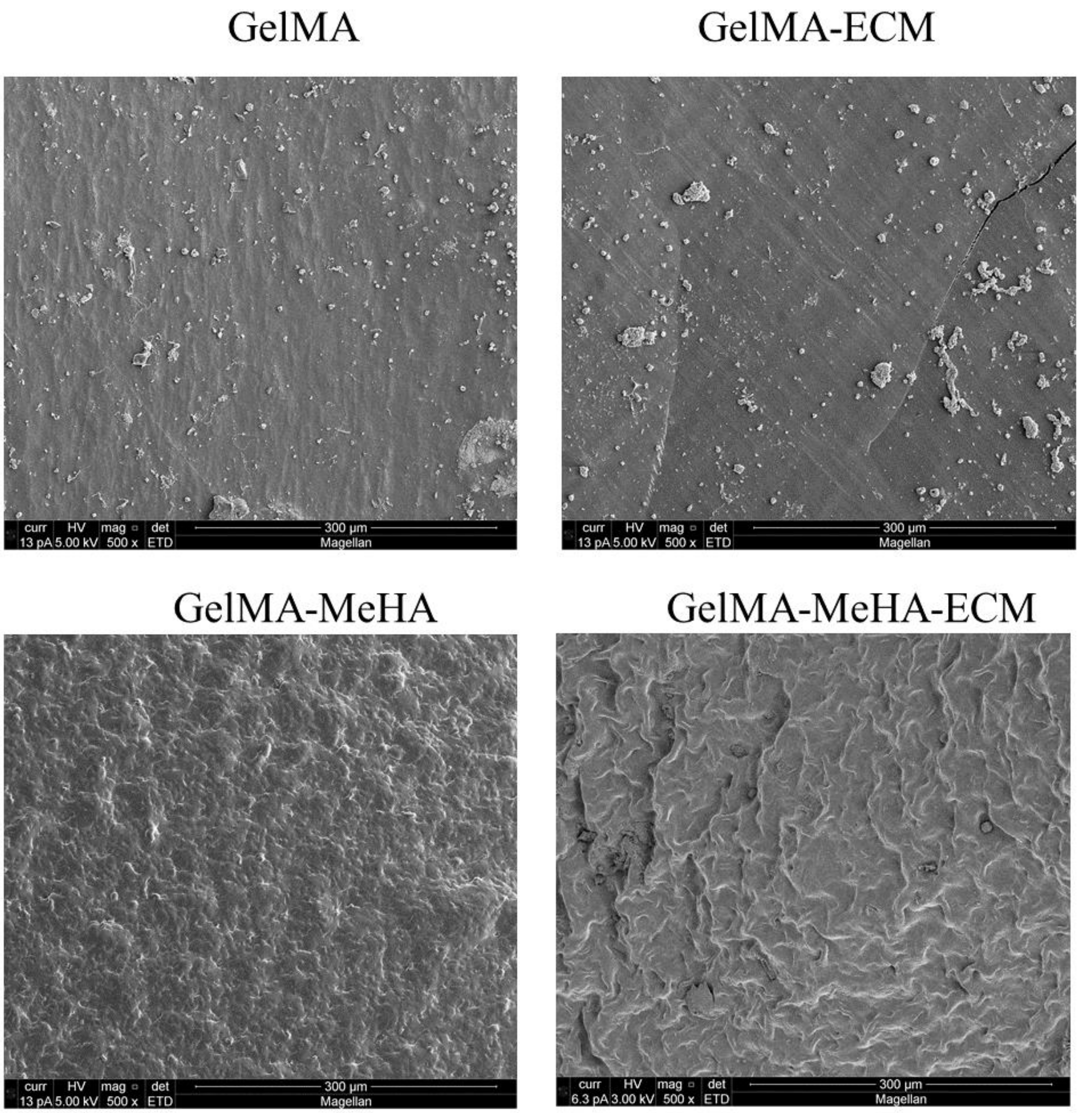


**Figure S1.** SEM images of GelMA, GelMA-dhECM, GelMA-MeHA, GelMA-MeHA-dhECM hydrogels.

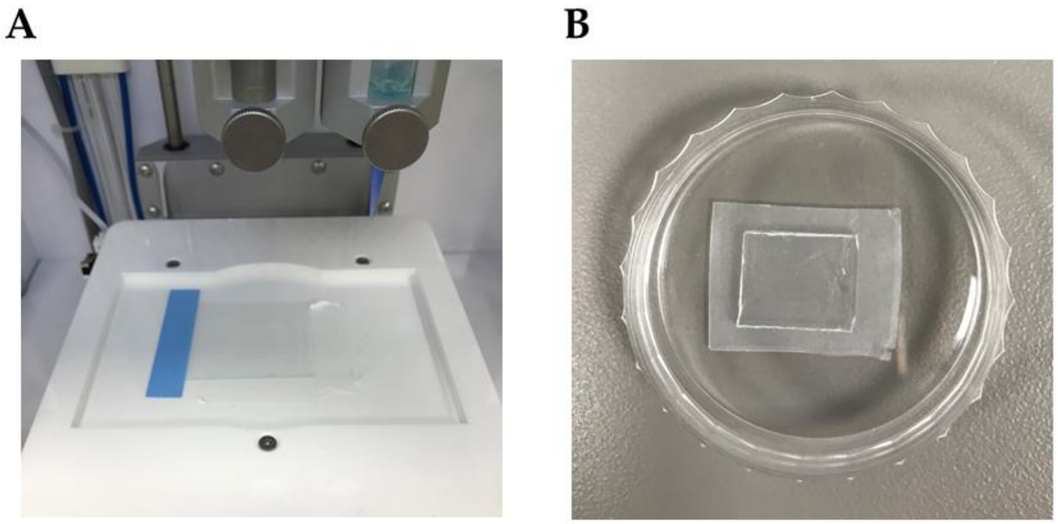

**Figure S2.** Parafilm coating (A) on the printing stage, (B) in the dish under the glass

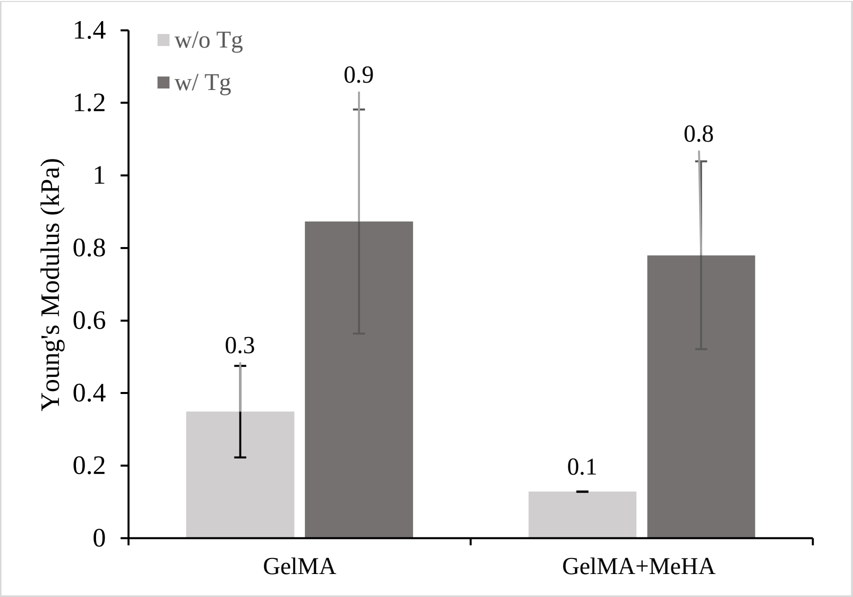


**Figure S3.** Young's Modulus of GelMA and GelMA-MeHA hydrogels with visible light crosslinking with and without mTGase treatment.

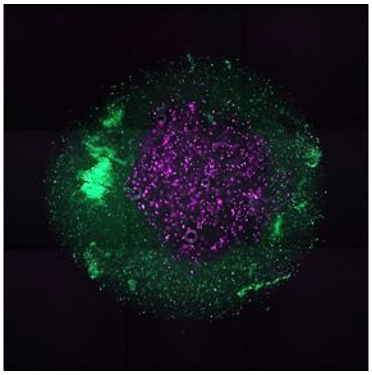

**Figure S4.** Printed boundary region using iCMs in GelMA-ECM bioink (green), and hCFs encapsulated in GelMA-MeHA-ECM bioink (magenta)
